# The RNA binding protein Nab2 genetically interacts with multiple RNA exosome cofactors to regulate target RNAs

**DOI:** 10.1101/2022.02.22.481433

**Authors:** Christy E. Kinney, Katherine Mills-Lujan, Milo B. Fasken, Anita H. Corbett

**Affiliations:** Department of Biology, and Graduate Program in Department of Human Genetics, Emory University, Atlanta, GA 30322 USA; Genetics and Molecular Biology, Department of Human Genetics, Emory University, Atlanta, GA 30322 USA; Department of Human Genetics, Emory University, Atlanta, GA 30322 USA

**Keywords:** Nab2, RNA binding protein, RNA exosome

## Abstract

RNA binding proteins play important roles in the processing and precise regulation of RNAs. Highlighting the biological importance of RNA binding proteins is the increasing number of human diseases that result from mutations in genes that encode these proteins. We recently discovered that mutations in the *ZC3H14* gene, which encodes an evolutionarily conserved polyadenosine RNA-binding protein, cause intellectual disability. Studies of the budding yeast orthologue of ZC3H14, Nuclear Poly(A) Binding protein 2 (Nab2), have provided insight into the functions of this protein. The *NAB2* gene is essential in *S. cerevisiae*, and conditional *nab2* mutants cause defects in a number of steps in RNA processing. To explore the critical functions of the Nab2/ZC3H14 protein family, we performed a high-copy suppressor screen on *nab2* mutant cells. This screen identified genes encoding two core subunits of the RNA exosome, as well as Nrd1 and Ski7, nuclear and cytoplasmic cofactors of the RNA exosome, respectively. Nrd1 is an RNA binding protein that is part of the Nrd1-Nab3-Sen1 (NNS) complex, which plays an important role in transcription termination of non-coding RNAs. Ski7 is a GTP-binding protein that mediates interaction between the RNA exosome and the Ski complex, which targets RNA transcripts to the exosome for processing and degradation in the cytoplasm. To explore the functional interactions between the RNA exosome and Nab2, we employed RNA-seq analysis to identify the transcripts most impacted by overexpression of these exosome cofactors in *nab2* mutant cells. This analysis revealed that many transcripts show small changes in steady-state levels, consistent with a global role of Nab2 in modulating transcript stability. This study uncovers functional interactions between the RNA exosome and Nab2 in both the nucleus and the cytoplasm.

## Introduction

RNA binding proteins comprise a diverse set of proteins critical for many aspects of post-transcriptional control of gene expression both in the nucleus and the cytoplasm. These proteins mediate key nuclear processing events, facilitate RNA export from the nucleus, regulate translation, and perform many other equally crucial cellular functions (1). Consistent with the numerous functions played by this class of protein, individual RNA binding proteins often contribute to multiple post-transcriptional regulatory steps required to fine tune gene expression. The importance of RNA binding proteins for proper cellular function is evident in the fact that many of these proteins are evolutionarily conserved (2) and the growing number of genes that encode RNA binding proteins that have been linked to human disease (3–6). Due to the evolutionary conservation of RNA binding proteins, a variety of model systems can be deployed to define their function and understand their multi-faceted roles in gene expression. An example of an evolutionarily conserved, multi-functional RNA binding protein linked to human disease is the zinc finger polyadenosine (polyA) RNA binding protein termed ZC3H14 in humans and Nab2 in budding yeast. Recent studies have identified mutations in *ZC3H14* that cause a non-syndromic autosomal recessive form of intellectual disability (5). Mutations in the essential *NAB2* gene in budding yeast cause defects in mRNA splicing (7), nuclear accumulation of poly(A) RNA (8, 9), and extended poly(A) tails (10). Rapid depletion of Nab2 from the nucleus causes a striking decrease in the level of polyadenylated RNAs (11, 12). Nab2, which is primarily localized to the nucleus, but can shuttle between the cytoplasm and the nucleus (13), binds with high affinity to polyadenosine RNA through a zinc finger poly(A) RNA-binding domain (10,14,15). How this polyadenosine RNA binding protein regulates multiple aspects of gene expression at the mechanistic level is not yet understood.

We exploited a yeast genetics approach to identify the most critical cellular functions of the Nab2 protein. For this work, we utilized a cold-sensitive mutant of *NAB2*, *nab2-C437S* (14–16). This amino acid substitution, which is located in the poly(A) RNA binding domain, significantly reduces binding of Nab2 to target RNAs (14, 15). A high-copy suppressor screen was performed with *nab2-C437S* cells to identify genetic interactors of Nab2. This screen identified both core components of the RNA exosome and nuclear and cytoplasmic RNA exosome cofactors (17). All subunits of the RNA exosome were subsequently tested as suppressors, revealing the specificity for suppression by *RRP41* and *RRP42*. As detailed by Mills-Lujan et al., a likely mechanism for suppression is impairment of RNA exosome function by destabilization of formation of the complex upon overexpression of Rrp41 or Rrp42 (17). Due to the conservation of these two subunits as far back as the archaeal exosome (18), the stoichiometry of the complex may rely on their precise regulation.

To test the hypothesis that impairment rather than enhancement of RNA exosome function suppresses the cold-sensitive growth phenotype, exosome mutants (19) were overexpressed in *nab2-C437S* cells (17). These variants contain mutations that impede procession of RNA through the central channel of the RNA exosome (19). When overexpressed, these exosome variants suppress the *nab2-C437S* growth phenotype similarly to overexpression of *RRP41* or *RRP42,* supporting the RNA exosome impairment hypothesis. The specificity of exosome subunits in this rescue of growth indicates the importance of the RNA exosome in mediating the growth of Nab2 mutant cells, potentially pointing to a key genetic interaction between the complex and Nab2.

While elucidating the mechanism of suppression by subunits of the RNA exosome, the previous study did not delve into the mechanism of suppression by the RNA exosome cofactors also identified in the screen. Here we focus on exploring how the overexpression of RNA exosome cofactors can suppress the cold sensitive growth of *nab2* mutant cells, providing insight into both the function of the evolutionarily conserved Nab2 protein and the cofactors that regulate RNA exosome function. Our results identify a link to control of the expression of ribosomal subunit genes that is mediated by the RNA exosome and nuclear RNA exosome cofactors but not a cytoplasmic cofactor, suggesting that multiple mechanisms link the RNA exosome and Nab2 function.

## Materials and Methods

### Saccharomyces cerevisiae strains and plasmids

Wildtype strain ACY233 (W303) and integrated mutant *nab2-C437S* strain ACY1026 (W303 background) were used in this study. Plasmid cloning was performed using the NEB Hifi assembly kit per manufacturer’s instructions. A 2µ *URA3* vector was employed for overexpression of high copy suppressor genes. Myc-tagged plasmids were constructed using empty 2µ *URA3* vector pAC3007 containing 2x-Myc and *ADH1* terminator sequences as the template.

For domain analysis, various deletions and amino acid substitutions in Nrd1 and Ski7 were created based on structural information available (20, 21).

### Saccharomyces cerevisiae transformations and growth assays

Yeast cells were grown overnight at 30°C to saturation in minimal media lacking uracil. Cell concentrations were measured and normalized to OD_600_= 0.4 and serially diluted in 10-fold dilutions. Cells were mixed and spotted onto minimal media plates lacking uracil and incubated at 18°C, 25°C, 30°C, and 37°C. Cells were also serially diluted and spotted on YPD plates for controls. Plates incubated at 25°C, 30°C, and 37°C were imaged after one and two days. Plates incubated at 25°C were also imaged after three days of growth. The plates grown at 18°C were imaged after three, four, and five days.

### Total RNA isolation

RNA was isolated from 2 mL liquid cultures grown overnight at 30°C to saturation, diluted into 10 mL cultures at OD_600_ = 0.4, and then grown at 25°C for 4 hours. Cultures were then spun down at 3,000 x g and resuspended in 1 mL TRIzol (Invitrogen). Cells were disrupted with glass beads using the Biospec Mini Bead Beater 16 Cell Disrupter at 25°C for 1 min, placed back on ice, and then disrupted again for 1 min. 100 µL of 1-bromo-3-chloropropane was added to each sample, followed by 15 sec on the vortex and a 2 min incubation at 25°C. Samples were then centrifuged at 16,000 x g at 4°C for 8 min. The upper layer of each sample was transferred to a fresh microcentrifuge tube, precipitated with 500 µL isopropanol, and then vortexed for 10 sec. RNA was centrifuged at 16,000 x g at 4°C for 8 min, and the pellet was washed with 1 mL of 75% ethanol. Each sample was centrifuged again at 16,000 x g at 4°C for 5 min and then air-dried. RNA was resuspended in 50 µL diethylpyrocarbonate [DEPC (Sigma)]-treated water to either be DNase-treated or frozen at −80°C. 2.5 µg of RNA from each sample was added to 2.5 µL amplification-grade DNase-I (Invitrogen), 2.5 µL DNase-I Reaction Buffer, and DEPC-treated water up to a volume of 25 µL. The reactions were incubated at 25°C for 15 min, followed by inactivation of DNase-I by the addition of 2.5 µL of EDTA 25 µM solution. Reactions were incubated at 65°C for 10 min and either frozen at −80°C or converted to cDNA.

### cDNA preparation and quantitative RT-PCR

1 µg DNase-treated RNA from each sample was reverse-transcribed to cDNA using M-MLV Reverse Transcriptase (Invitrogen) according to the manufacturer’s protocol. 1 µg DNase-treated RNA from each sample was simultaneously PCR amplified without M-MLV Reverse Transcriptase to be used as a control. For quantitative PCR, 10 ng of each sample (technical triplicates of 3 independent biological replicates) was amplified using various primers (0.5 µM; Table S2) and SYBR Green PCR master mix (QIAGEN). Reactions were run for 44 cycles on a StepOnePlus Real-Time PCR machine (Applied Biosystems) with an annealing temp of 55°C. Statistical means were calculated and compared using the 2^-ΔΔCt^ method (22). Mean RNA levels from experimental samples were normalized to mean RNA levels from wildtype samples, and normalized means were graphed as relative fold change of experimental samples to wildtype with error bars representing standard deviation.

### RNA-sequencing and computational analysis

2 µg total RNA from each sample were analyzed for quality (Bioanalyzer), depleted of rRNA (Illumina RiboZero Gold kit), and prepared into reverse-stranded libraries (Roche KAPA kit). RNA samples were sequenced on the NextSeq PE75 High-Output Flow Cell platform. Biological duplicates were prepared and processed for all conditions. Paired-end raw reads were concatenated using Galaxy (23). Reads were aligned to *Saccharomyces cerevisiae* S288C genome assembly R64.1.1 (SGD) using STAR (24), and counted using featureCounts (25). Gene annotations were downloaded from SGD, and CUTs and SUTs were also annotated (26). Raw read counts were rlog normalized for principal component analysis (PCA) and hierarchical clustering. Differential expression analysis was performed using the DESeq2 package (R V 3.6.2, (27)). Differential expression results were calculated based on gene expression of experimental samples compared to gene expression of WT samples or gene expression of nab2 mutant samples. Gene ontology was performed using Gene Set Enrichment Analysis (GSEA 4.1.0) (28, 29). Samples were run using a pre-ranked gene list generated from the DESeq2 output. Adjusted p-values of genes were normalized using log base 10 and ranked from highest to lowest. We ran 1000 permutations and excluded gene sets with >500 or <15 genes per GSEA default.

### Data visualization

R (v3.6.2) was used to generate figures from normalized read counts or reads from differential expression analysis. Figures were built with the following R packages: ggplot2 (30), gplots (31), and RColorBrewer (32).

## Results

### Specific RNA exosome cofactors suppress the growth defect of *nab2-C437S* cells

A previous screen for high-copy suppressors of the *nab2-C437S* cold-sensitive phenotype (17) identified RNA exosome cofactors as putative high copy suppressors. The RNA exosome is a conserved ribonuclease complex with both exonuclease and endonuclease activities (19,33–35). It is localized to the nucleus, nucleolus, and cytoplasm, functioning in RNA processing and quality control pathways. RNA exosome cofactors assist the exosome by directing it to particular RNA targets to perform specific functions. To test whether RNA exosome cofactors can suppress *nab2-C437S* cold-sensitive growth defect, we performed a 10-fold serial dilution growth assay (Figure 1A). *nab2-C437S* cells were transformed with vector alone or, as a control, *NAB2*, as well as RNA exosome subunit genes *RRP41* and *RRP42*, which have been validated as suppressors of *nab2-C437S* (17). Figure 1A shows that the nuclear RNA exosome cofactors Nrd1, a component of the Nab3-Nrd1-Sen1 (NNS) complex which functions in transcription termination of short, primarily non-coding RNAs (36), and Ski7, which recruits the Ski complex to the RNA exosome to target aberrantly processed mRNAs for degradation and to assist in mRNA turnover (37, 38), both suppress the *nab2-C437S* growth phenotype. Nucleolar RNA exosome cofactor Nop8, which participates in processing of 5.8S pre-rRNA and, ultimately, biogenesis of the 60S ribosome (39, 40), was also identified as a suppressor.

**Figure 1:**
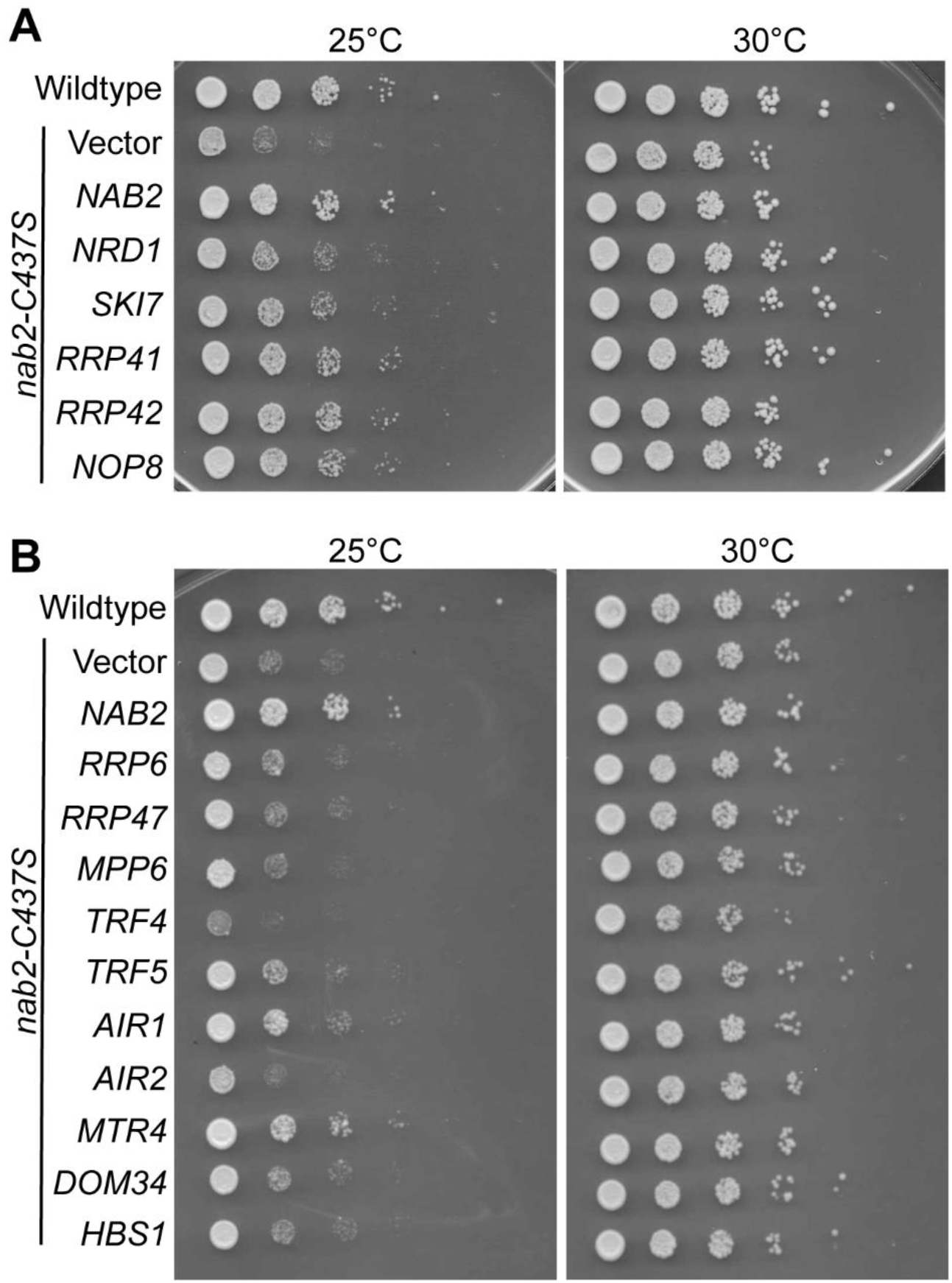
Suppression of the *nab2-C437S* cold-sensitive growth defect is specific to RNA exosome cofactors *SKI7*, *NRD1*, *NOP8*, and *MTR4*. *Nab2-C437S* cells show a cold-sensitive growth defect at 25°C. Suppressor genes were identified through a high-copy suppressor screen. (A) *nab2-C437S* cells were transformed with 2µ high-copy plasmids containing empty vector or genes identified in the high-copy suppressor screen. Cells were then serially diluted, spotted onto solid media plates, and grown at either 25°C or 30°C for 1-2 days. Overexpression of RNA exosome cofactor genes *NRD1*, *SKI7*, and *NOP8*, as well as RNA exosome subunit genes *RRP41* and *RRP42* suppress the cold-sensitive growth defect. (B) To test for specificity of RNA exosome cofactors as suppressors of the *nab2-C437S* growth defect, cells were transformed with 2µ plasmids containing nuclear and cytoplasmic cofactor genes. Cells were then serially diluted spotted onto solid media plates and grown at either 25°C or 30°C for 1-2 days.

To assess whether suppression of *nab2* growth defects is a general property of RNA exosome cofactors, we overexpressed a panel of RNA exosome cofactors in *nab2-C437S* cells (Figure 1B). Nuclear cofactors include the exonuclease Rrp6 and its interacting partner Rrp47, as well as Mpp6, a cofactor involved in RNA surveillance (41). We also overexpressed genes encoding members of the nuclear TRAMP complex: Trf4, Trf5, Air1, Air2, and Mtr4. The TRAMP complex is responsible for polyadenylation of transcripts targeted for degradation by the RNA exosome (42, 43). We also overexpressed cytoplasmic RNA exosome cofactors Dom34 and Hbs1, which comprise the Dom34-Hbs1 ribosome dissociation complex that releases stalled ribosomes from aberrantly processed mRNAs (44–46). In addition to cofactors Nrd1, Ski7, and Nop8, we also identified Mtr4 as a robust suppressor of the cold-sensitive growth phenotype of *nab2-C437S* cells. This distinct subset of RNA exosome cofactors that serve as suppressors highlights the specificity of interactors.

### Suppression of the *nab2-C437S* cold-sensitive growth phenotype is specific to the Nrd1 subunit of the NNS complex

Nrd1 functions primarily as part of the nuclear Nab3-Nrd1-Sen1 (NNS) complex in transcription termination of short RNAs. Nrd1 and Nab3 are essential RNA binding proteins, while Sen1 is a DNA/RNA and RNA helicase. The complex co-transcriptionally interacts with Polymerase II to target transcripts for transcription termination or for degradation in coordination with the TRAMP complex and the RNA exosome. Outside of the functions in the NNS complex, Nrd1 and Nab3 are also recruited to misprocessed mRNAs in a quality control pathway and can bind independently to sites within the genome, suggesting these proteins could have independent functions (47, 48). To determine if suppression of the *nab2-C437S* cold-sensitive growth phenotype is Nrd1-specific, we performed a growth assay and overexpressed each NNS component in *nab2-C437S* cells. Neither overexpression of *NAB3* nor *SEN1* suppressed the *nab2-C437S* growth phenotype (Figure 2A). The specificity of Nrd1 as an independent suppressor of the *nab2-C437S* cold-sensitive growth phenotype, suggests a possible role for Nrd1 outside of the NNS complex.

**Figure 2:**
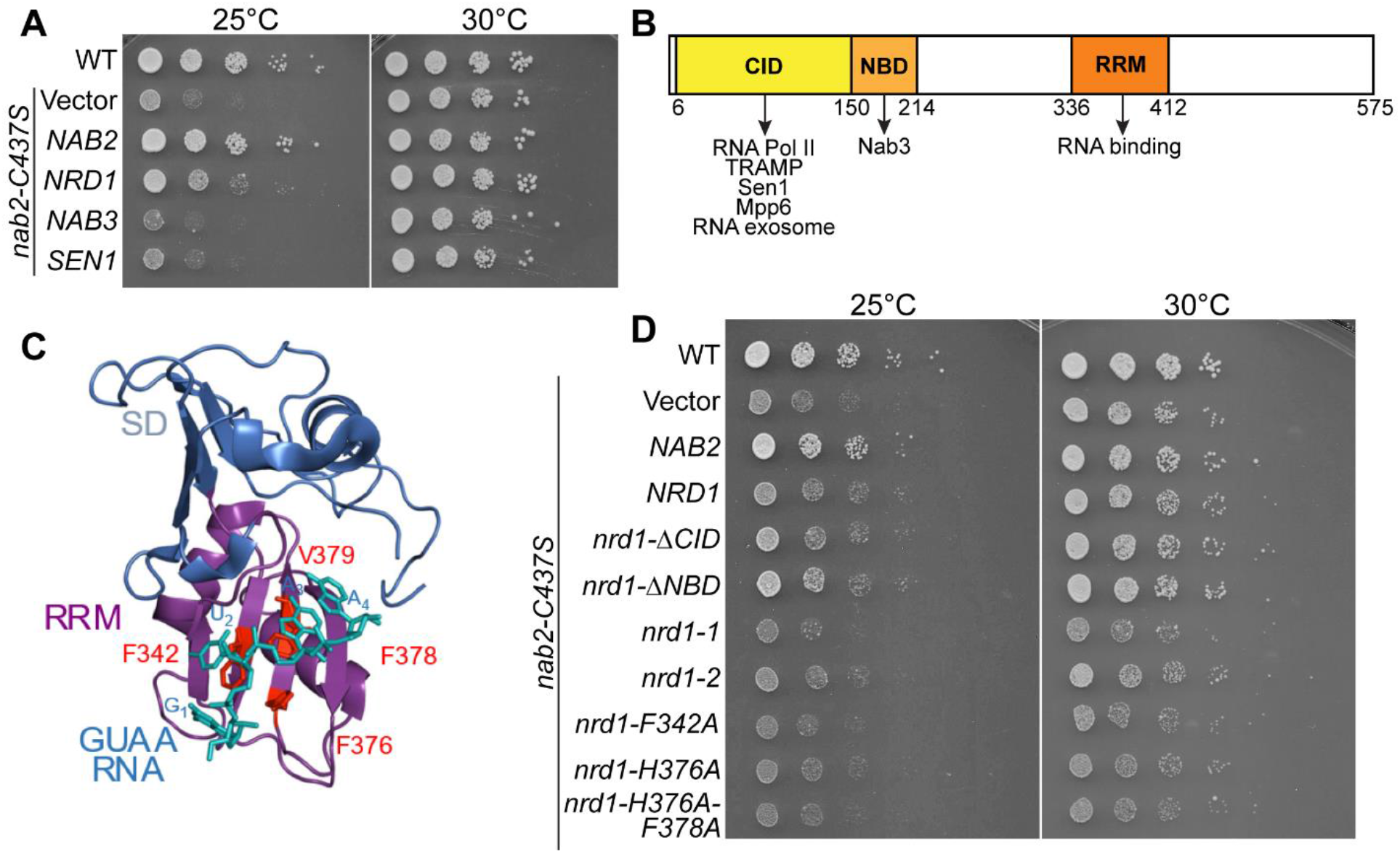
Suppression is specific to *NRD1* of the Nrd1-Nab3-Sen1 (NNS) complex and requires the Nrd1 RNA binding domain. (A) To test for specificity of suppression by *NRD1*, 2µ plasmids containing NNS subunit genes *NRD1*, *NAB3*, and *SEN1* were transformed into *nab2-C437S* cells. Cells were serially diluted, spotted onto solid media plates, and grown at 25°C or 30°C for 1-2 days. (B) Nrd1 domain structure with domain-specific interactions indicated by arrows. CID, Polymerase II C-Terminal Interacting Domain; NBD, Nab3 Binding Domain; RRM, RNA Recognition Motif. (C) Crystal structure of the essential Nrd1 RRM and interacting GUAA RNA (20). Indicated amino acids are required for RNA binding, and Nrd1 variants with impaired RNA binding function were generated by changing these amino acids. (D) To test for Nrd1 domains that are required for suppression, *nab2-C437S* cells were transformed with 2µ plasmids containing *NRD1* mutants lacking the CID (*ΔCID*), lacking the NBD (*ΔNBD*), or with impaired RNA binding (premature nonsense mutants *nrd1-1*, *nrd1-2*; *nrd1-F342A*, *nrd1-H376A*, *nrd1-H376A-F378A*).

### The RNA binding function of Nrd1 is required for *nab2-C437S* suppression

As illustrated in Figure 2B, Nrd1 contains three functionally important domains: the RNA polymerase II C-terminal interacting domain (CID); the Nab3 binding domain (NBD), and the RNA recognition motif (RRM). The Nrd1 CID interacts in a mutually exclusive manner with either the C-terminal domain of Polymerase II or with Trf4 of the TRAMP complex. This domain is also responsible for direct interactions between Nrd1 and the NNS partner Sen1 as well as with RNA exosome cofactors Rrp6 and Mpp6. The NBD mediates interactions with Nab3, while the RRM mediates RNA binding to GUA[A/G] termination elements of RNAs, as well as other G-rich and AU-rich sequences (20,36,49,50).

We performed a serial dilution spotting assay to determine which functional domains and interactions of Nrd1 are required for suppression of the cold-sensitive growth phenotype of *nab2-C437S* cells. Although Nrd1 is essential, both the CID and NBD domains can be deleted and the protein remains functional (49). Thus, we generated Nrd1 deletion variants for the CID and the NBD. The Nrd1 RRM is essential (49). However, structural studies have defined critical residues within the RRM that mediate RNA binding (Fig 2C) and subsequent biochemical studies were performed to demonstrate that specific residues (F342A, H376A, and F378A) within the Nrd1 RRM are required for high affinity binding to RNA (51). Thus, we could take advantage of this information to generate characterized Nrd1 variants that disrupt the Nrd1/Pol II interaction and the Nrd1 interaction with RNA. As shown in Figure 2D, the Nrd1 ΔCID and ΔNBD variants both suppressed the growth phenotype of *nab2-C437S* cells, while each of the Nrd1 RRM variants failed to suppress (Figure 2D). This result indicates that the interaction of Nrd1 with target RNAs specifically is required to suppress the *nab2* growth defect. The finding that the Nab3 binding domain of Nrd1 is dispensable for suppression further bolsters the idea that this suppression could be mediated in a Nab3-independent manner.

### Suppression of the *nab2-C437S* cold-sensitive growth phenotype is specific to Ski7 and not conferred by other subunits of the Ski complex

In the cytoplasm, the RNA exosome works in conjunction with the Ski complex in a general mRNA decay pathway, as well as to degrade non-stop RNAs (33,52,53). The Ski complex is a tetramer comprised of the helicase Ski2, the tetratricopeptide protein Ski3, and two copies of the WD repeat protein Ski8 (54). The interaction between these two complexes is mediated by Ski7. To test whether the suppression of cold-sensitive growth is specific to *SKI7* overexpression, we overexpressed the genes encoding each of the 3 different subunits of the Ski complex in *nab2-C437S* cells. While overexpression of *SKI7* showed clear suppression, overexpression of *SKI2*, *SKI3*, and *SKI8* did not rescue *nab2-C437S* growth (Figure 3A).

**Figure 3:**
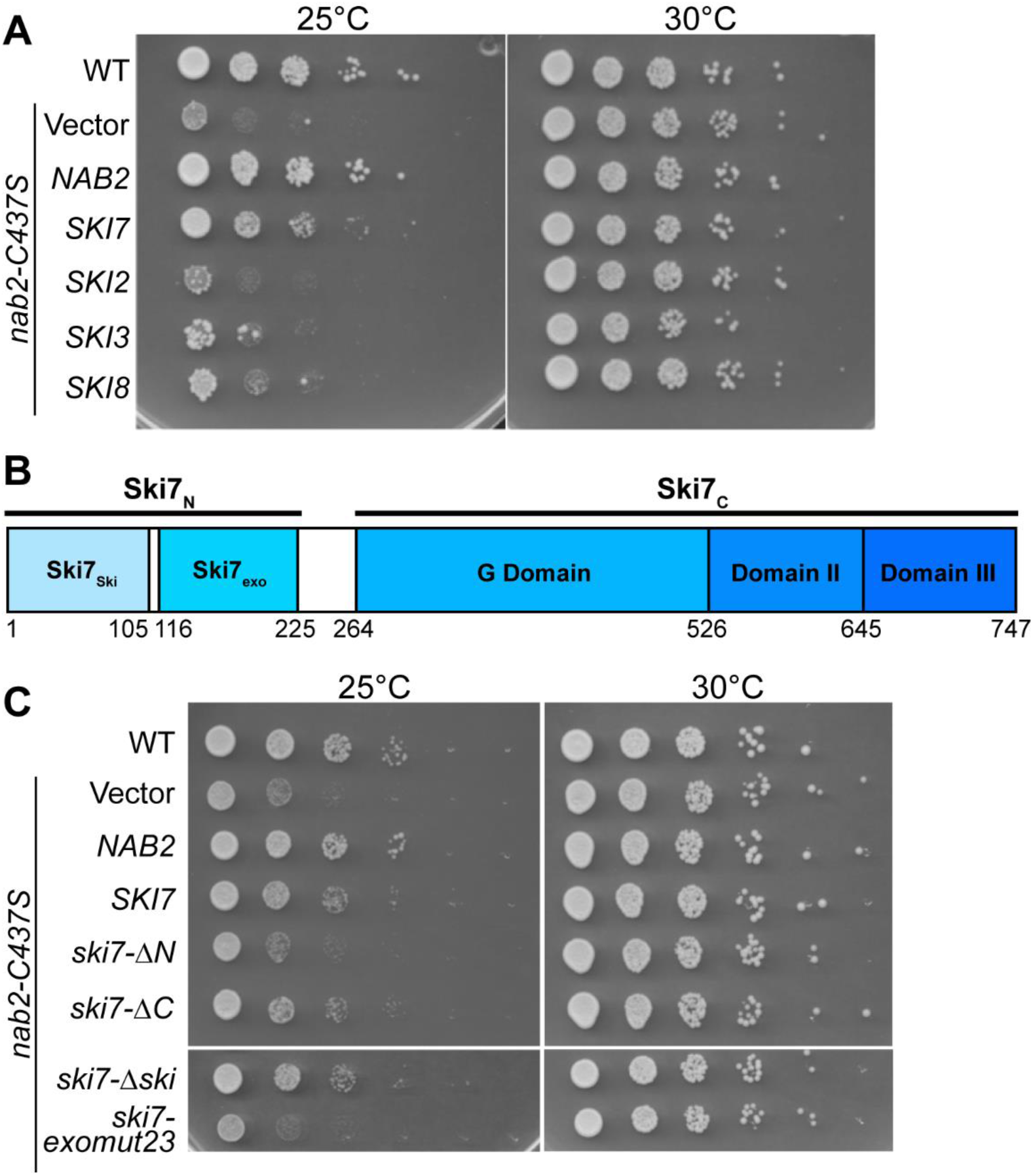
Suppression is specific to *SKI7* and requires the Ski7-RNA exosome interacting domain. (A) *nab2-C437S* cells were transformed with 2µ plasmids containing subunits of the cytoplasmic RNA exosome cofactor Ski complex. Cells were serially diluted, spotted onto solid media plates, and grown at 25°C or 30°C for 1-2 days. (B) Ski7 domain structure. Ski7_ski_, Ski complex-interacting domain; Ski7_exo_, RNA exosome-interacting domain. (C) *nab2-C437S* cells were transformed with 2µ plasmids containing SKI7 mutants lacking the C-terminal domain (*ΔC*), lacking the N-terminal domain (*ΔN*), lacking the N-terminal Ski-complex interacting domain (*Δski*), or mutations in the N-terminal RNA exosome interacting domain that interrupt binding to the RNA exosome (*exomut23*).

### The interaction between Ski7 and the RNA exosome is required for suppression of the *nab2-C437S* growth phenotype

Ski7 has two functionally important domains, which include the N-terminal domain and the C-terminal domain (Figure 3B). The N-terminal domain is critical for interactions between Ski7 and both the ski complex and exosome complex (37,38,55,56). These interactions occur in distinct regions of the N-terminal domain. The C-terminal domain sequence encodes a pseudo-GTPase with a proposed role in degradation of non-stop RNAs (56). To determine which domains of Ski7 are required for suppression of the *nab2* growth phenotype, we generated Ski7 variants consisting of either the N-terminal RNA exosome interacting domain or the C-terminal GTP-binding domain. We then generated variants of the N-terminal domain consisting of either the functional Ski complex-interacting domain or the functional exosome-interacting domain based on structural studies of Ski7 (21, 38). The Ski7 variant with a functional Ski complex-interacting domain (*ski7-exomut23*) combines amino acid changes in two of four patches in the Ski7 N-terminal exosome-interacting domain (21). Disrupting the amino acid sequences of these interaction patches has been shown to eliminate binding between Ski7 and the exosome. The *ski7 Δski* variant, which retains a functional exosome-interacting domain, eliminates interaction between Ski7 and the Ski complex. We performed a serial dilution spotting assay and found that the N-terminal domain of Ski7 is required for suppression of the *nab2-C437S* cold-sensitive growth phenotype, indicating the requirement for interaction between Ski7 and the Ski complex, the exosome, or both (Figure 3C). Additionally, we found that the interaction between Ski7 and the exosome, but not the interaction between Ski7 and the Ski complex, is required for suppression of the *nab2-C437S* growth defect (Figure 3C).

### Overexpression of suppressors results in altered transcript profiles

RNA-seq analysis was performed to elucidate the transcripts and functional categories of transcripts affected in *nab2-C437S* variant cells, as well as the transcripts rescued by overexpression of suppressor genes *NRD1*, *SKI7*, *RRP41*, and *RRP42*. Principal component analysis on normalized transcript counts displays the tight clustering of biological replicates as well as the distinctness or relatedness of the different genotypes (Figure 4A).

**Figure 4:**
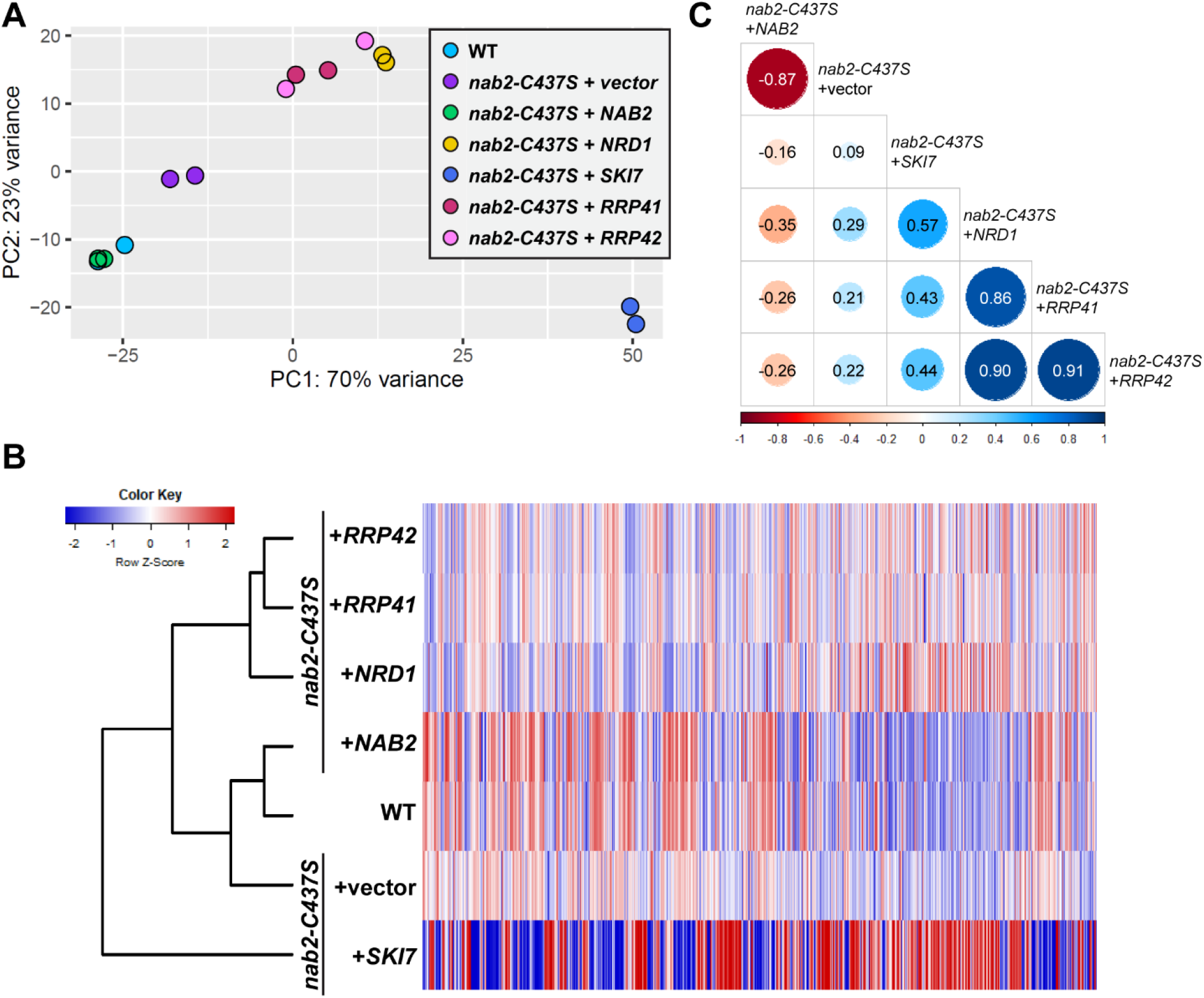
RNA-sequencing reveals distinct transcriptomes for *nab2-C437S* cells overexpressed with suppressor genes. RNA-sequencing was performed on biological duplicate samples of each genotype. (A) Principal component analysis (PCA) performed comparing rlog normalized gene counts for each biological replicate of all genotypes. (B) Heatmap showing rlog normalized gene count comparisons for combined biological replicates of each genotype. Heatmap is oriented horizontally. Each gene count is individually scaled across samples. (C) Correlation matrix showing correlations between differentially expressed genes of each genotype (combined replicates). Circle sizes correspond to degree of correlation. All correlation values are statistically significant, with sstatistical significance set at p<0.05.

Displayed in the heatmap (Figure 4B), hierarchal clustering of normalized counts for combined biological replicates also shows the relatedness of genotypes. The transcriptome of *nab2-C437S* cells was altered significantly from that of wildtype cells. Overexpression of *NAB2* in the *nab2-C437S* background resulted in a transcript profile most similar to wildtype control cells, suggesting possible restoration of steady-state levels of transcripts altered by Nab2-C437S. In contrast, overexpression of suppressor genes *NRD1*, *SKI7*, *RRP41*, and *RRP42* in *nab2-C437S* cells resulted in distinct transcript profiles compared to overexpression of *NAB2*. Overexpression of suppressors *RRP41* and *RRP42* resulted in extremely similar transcript profiles. This is not surprising, as both suppressor genes encode subunits of the RNA exosome. Overexpression of *NRD1* resulted in a fairly strong degree of transcript profile overlap with the *RRP41* and *RRP42* genotypes; however, as an RNA exosome cofactor rather than subunit, *NRD1* overexpression was not as closely clustered to the exosome subunit samples as the subunit overexpression samples were to each other. Overexpression of *SKI7* resulted in the most distinct transcript profile, clustering away from the other suppressors and displaying the most extreme steady-state level changes of transcripts of all samples. The effects of overexpression of cytoplasmic RNA exosome cofactor *SKI7* on RNAs in the cytoplasm may account for this distinct profile.

Differential expression analysis was subsequently performed on samples (Figure 4C). Samples were both compared to wildtype expression levels and to *nab2-C437S* expression levels. Like the normalized counts, the correlations between samples showed the highest degree of relatedness between the *RRP41* and *RRP42* overexpression samples. The *NRD1* overexpression sample closely correlated with the subunits, while *SKI7* was only moderately correlated to any other sample.

### The *nab2-C437S* mutation affects a small subset of gene ontology categories

*Nab2-C437S* cells have sets of transcripts with both increased and decreased steady-state levels (Figure 5A). The majority of affected transcripts showed a log_2_ fold change smaller than 1. To determine the gene ontology categories affected by *nab2-C437S*, Gene Set Enrichment Analysis (GSEA) was performed. This analysis revealed 14 gene ontology (GO) categories significantly decreased in *nab2-C437S* cells compared to control cells (Figure 5B). Of these GO terms, several overarching categories encapsulate multiple listed terms. These include ribosome subunits and biogenesis, polymerase activity, and telomere maintenance. Zero GO terms were significantly increased compared to wildtype cells. Overexpression of *NAB2* resulted in rescue of most significantly decreased GO categories in *nab2-C437S* cells (Figure 5C), highlighting the importance of Nab2 function in regulating these cellular processes. Of note, Nab2 regulates numerous ribosomal transcripts, suggesting a link to regulation of translation.

**Figure 5:**
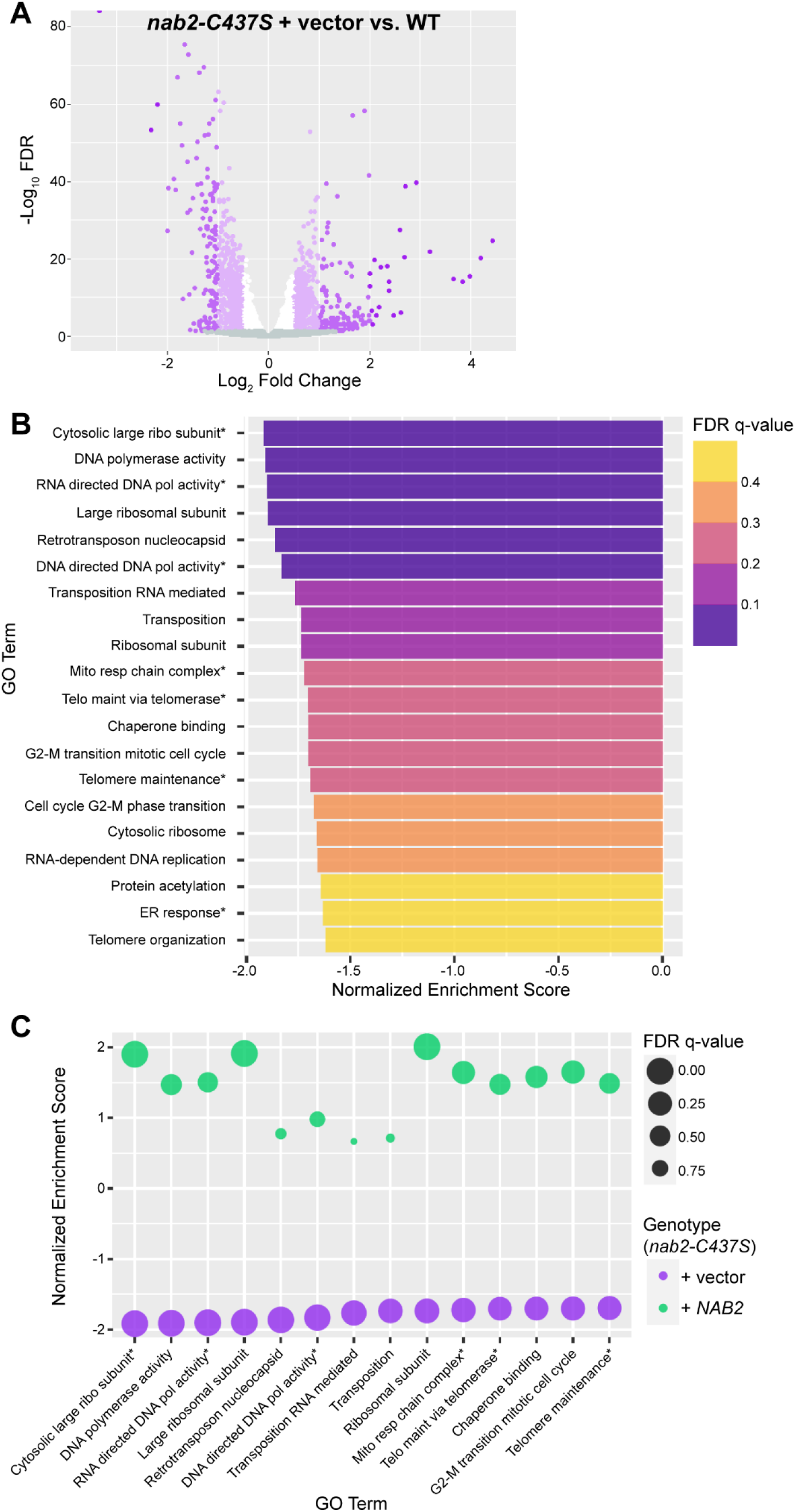
*nab2-C437S* cells show 14 gene ontology categories significantly down compared to control cells. (A) Volcano plot showing differentially expressed genes in *nab2-C437S* cells compared to control cells. (B) Bar chart showing the most decreased GO terms in *nab2-C437S* cells compared to control cells. Categories with an FDR q-value <0.25 (including dark purple, medium purple, and pink bars) were considered for further analyses. (C) Bubble plot showing comparison of the 14 GO categories most significantly down in *nab2-C437S* cells (compared to control cells) against *nab2-C437S* cells overexpressed with wildtype *NAB2* (compared to *nab2-C437S* cells). (B-C) Asterisk (*) used to indicate truncated category names and/or abbreviated terms. [ribo, ribosomal; pol, polymerase; telo, telomere; maint, maintenance] [Mito resp chain complex, Mitochondrial respiratory chain complex assembly; Telomere maintenance <u>via telomere lengthening</u>; ER response, Endoplasmic reticulum unfolded protein response & response to endoplasmic reticulum stress & cellular response to unfolded protein]

### Overexpression of suppressors results in significant overlap of affected transcripts and GO categories

Overexpression of suppressors *NRD1*, *SKI7*, and *RRP41* in *nab2-C437S* cells results in overlapping sets of affected transcripts among suppressors (Figure 7). Transcripts with significantly increased or decreased steady-state levels resulting from *RRP41* overexpression were largely and similarly altered by *NRD1* overexpression, showing an overlap of 89%. While the majority of transcripts affected by *RRP41* overexpression were also affected by *NRD1* overexpression, *NRD1* overexpression impacted a large number of transcripts unaffected by RRP41 overexpression. Transcripts affected by *NRD1* overexpression share a greater overlap with transcripts affected by *SKI7* overexpression than with transcripts affected by *RRP41* overexpression. The overlap of transcripts altered by overexpression of suppressor gene *RRP41*, which is both nuclear and cytoplasmic, with transcripts affected by overexpression of nuclear RNA exosome cofactor *NRD1* suggests that the most critical transcripts for suppression may be nuclear. Indeed, the category most GO category most affected in *nab2-C437S* cells, *Translation*, is rescued by both Nrd1 and Rrp41, suggesting that the Nrd1-mediated suppression is likely through regulation of genes in this category such as ribosomal subunit genes.

**Figure 6:**
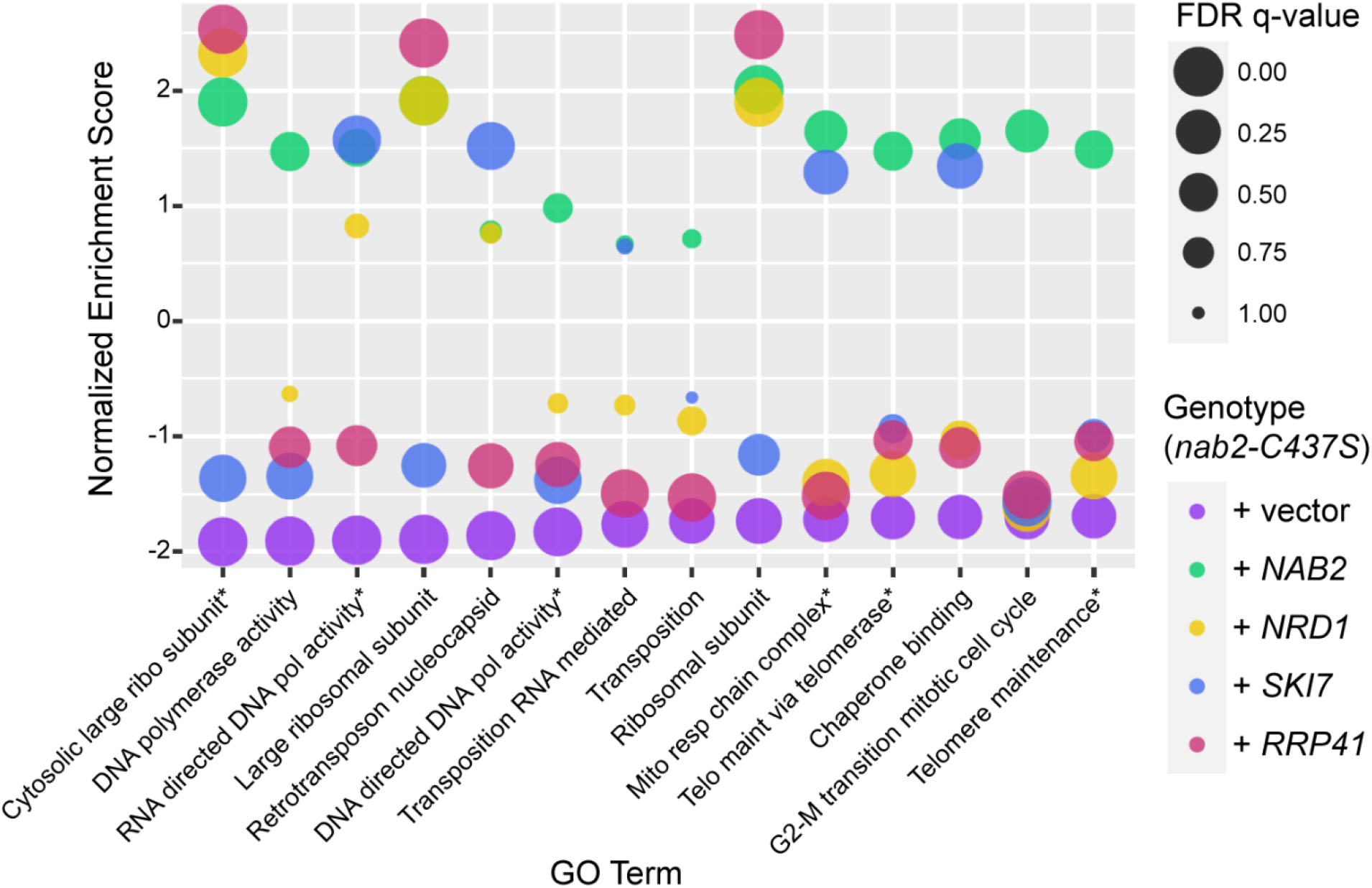
The top 14 down categories in *nab2-C437S* cells show disparate patterns of rescue/lack of rescue upon overexpression of suppressors *NRD1*, *SKI7*, and *RRP41*. Bubble plot showing comparison among samples for the 14 most significantly down categories identified in *nab2-C437S* cells. Asterisk (*) used to indicate truncated category names and/or abbreviated terms. [ribo, ribosomal; pol, polymerse; telo, telomere; maint, maintenance] [Mito resp chain complex, Mitochondrial respiratory chain complex assembly; Telomere maintenance <u>via telomere lengthening</u>]

**Figure 7:**
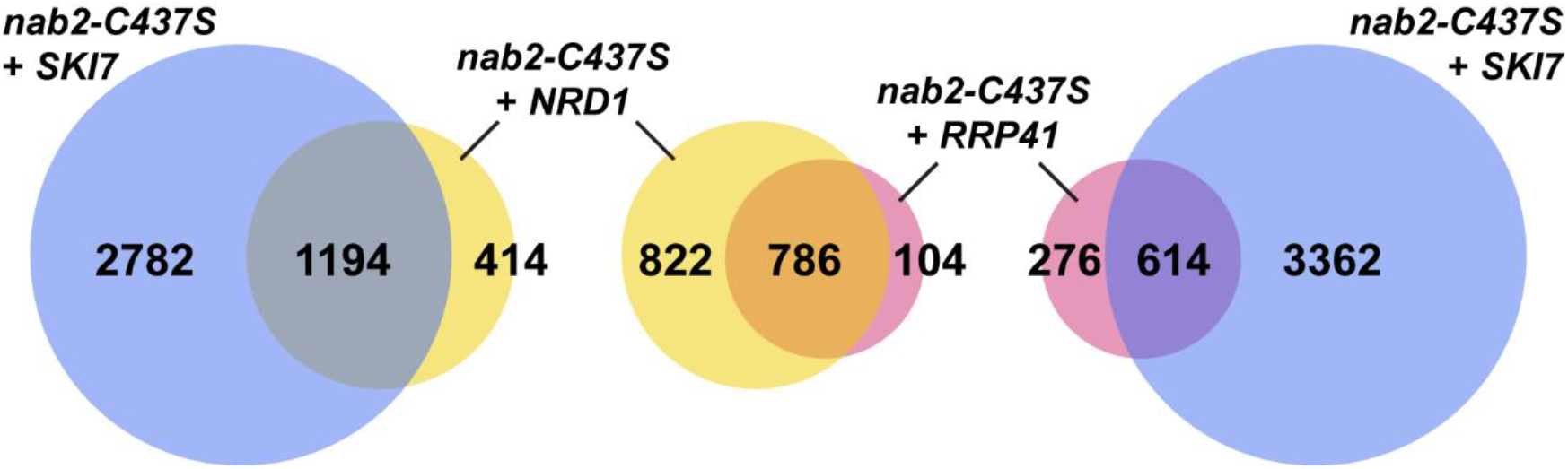
Overexpression of suppressors in *nab2-C437S* cells results in significant overlap of differentially expressed genes between different suppressor samples. Lists of differentially expressed genes for *nab2-C437S* cells overexpressed with *NRD1*, *SKI7*, or *RRP41* were compared for each pair of suppressors. Gene counts include transcripts with both up and down steady-state levels. Absolute log_2_ fold change cut-off set at α>0.5. Statistical significance set at p<0.05. Circles drawn to scale.

Following a similar trend of overlap as transcripts, Gene Set Enrichment Analysis showed overlap of GO categories among suppressor samples (Figure 8A). To determine the similarities in categories most affected by *NRD1* or *SKI7* overexpression, the top ten most positively and negatively enriched GO categories were compared among samples (Figure 8B,C). Of the categories most decreased by *NRD1* overexpression, all were similarly decreased by overexpression of *SKI7* or *RRP41*. Of the categories most increased by *NRD1* overexpression, all were similarly increased by overexpression of *RRP41*. Half of these increased categories were also increased upon *SKI7* overexpression; however, categories pertaining to the cytosol were decreased. Of the categories most decreased by *SKI7* overexpression, all were similarly decreased by *NRD1* overexpression. Notably, overexpression of *RRP41* resulted in an increase for nine out of ten of these categories. Of the categories most increased by *SKI7* overexpression, all categories were similarly increased by either *NRD1* or *RRP41* overexpression.

**Figure 8:**
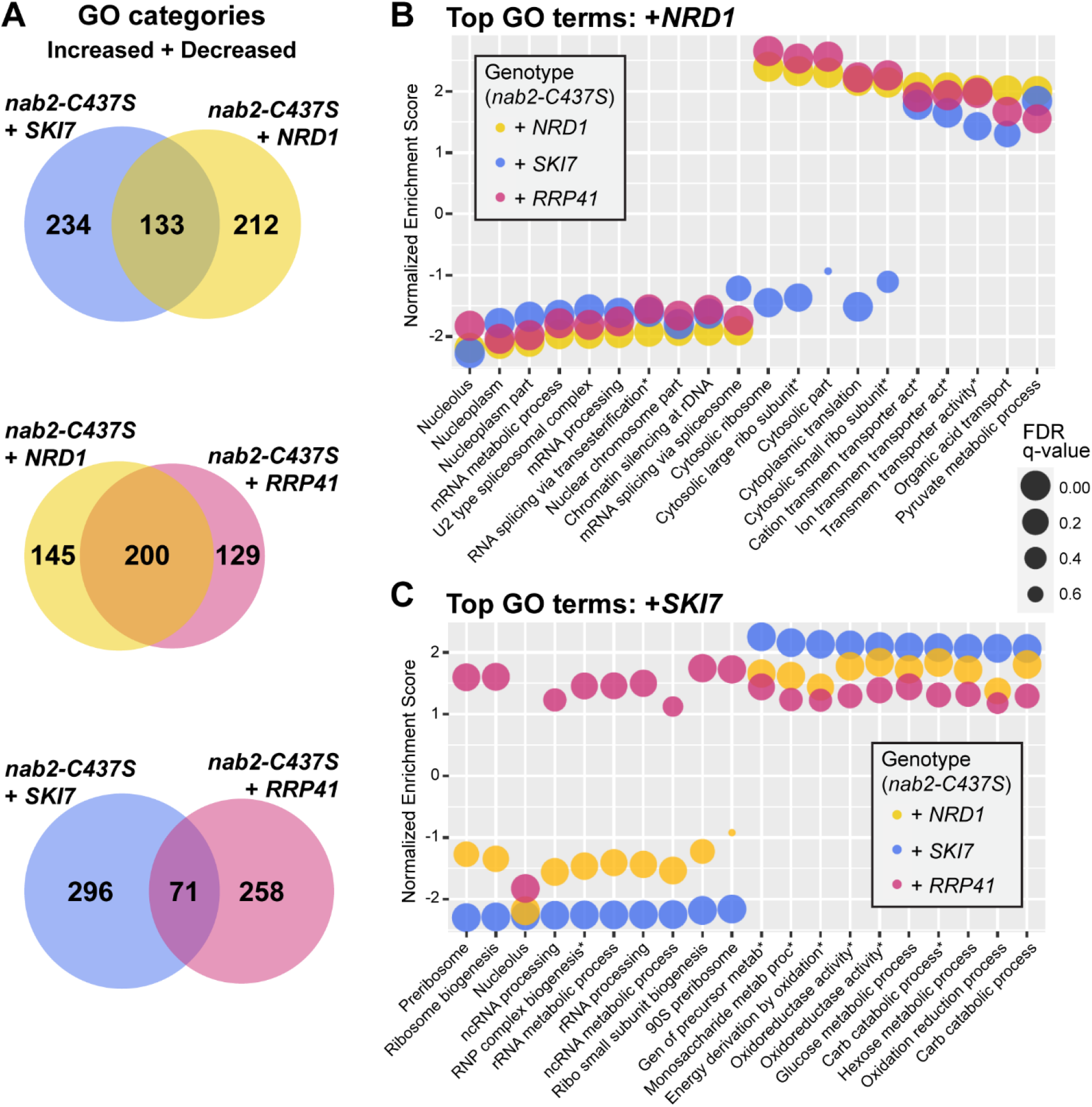
Overexpression of *NRD1*, *SKI7*, and *RRP41* results in significant overlap of differentially expressed gene ontology categories affected by each suppressor. (A) Venn diagrams showing comparisons of differentially expressed gene ontology categories comparing each pair of suppressors in *nab2-C437S* cells. Totals include both up and down gene ontology terms. Absolute log_2_ fold change cut-off set at α>0.5. Statistical significance set at p<0.05. Circles drawn to scale. (B) Bubble plot showing the top 10 down and top 10 up GO terms for *nab2-C437S* cells overexpressed with *NRD1*. Matching GO terms were then subset from *SKI7* and *RRP41* overexpression datasets and plotted. (C) Bubble plot showing the top 10 down and top 10 up GO terms for *nab2-C437S* cells overexpressed with *SKI7* with matching GO terms subset and plotted from *NRD1* and *RRP41* overexpression datasets. (B,C) Log_2_ fold change cut-offs set at α>0.5 or α<-0.5. Statistical significance set at p<0.05. [ribo, ribosomal; transmem, transmembrane; act, activity; RNP, ribonucleoprotein; carb, carbohydrate] [Gen of precursor metab, Generation of precursor metabolites and energy; Monosaccharide metab proc, Monosaccharide metabolic process; Energy derivation by oxidation <u>of organic compounds</u>; Oxidoreductase activity <u>acting on CH OH group of donors</u>; Oxidoreductase activity <u>acting on the CH OH group of donors NAD or NADP as acceptor</u>]

### Transcript levels altered by overexpression of suppressors are not altered in wildtype control cells

Overexpression of *NRD1*, *SKI7*, or *RRP41* alters the steady-state levels of thousands of genes in *nab2-C437S cells*. As elucidating the functions and interactions of Nab2 was the main goal of this project, a concern arose that, rather than shedding light on Nab2 function, the RNA-seq data was revealing how RNA exosome impairment independently affects the transcriptome. To determine the extent to which overexpression of cofactors and subunits affected the transcriptome independent of Nab2, we performed RNA-sequencing on samples overexpressing *NRD1*, *SKI7*, or *RRP41* in wildtype control cells. Surprisingly, we found significantly fewer affected transcripts in control cells and little overlap of altered transcripts between samples analyzed (Figure 9). *NRD1* overexpression in the control cells significantly altered the steady-state levels of 13 transcripts, compared to almost 4000 affected transcripts in *nab2-C437S* cells. *SKI7* overexpression increased the steady-state levels of 149 transcripts, while only five show decreased steady-state levels in control cells. Finally, *RRP41* overexpression affected the steady-state levels of 273 transcripts in the control cells, the greatest number amongst the suppressors, but this number is less than 10% of the total number of transcripts increased or decreased in the *nab2-C437S* cells. Of the 273 transcripts, only 63 were affected in both backgrounds.

**Figure 9:**
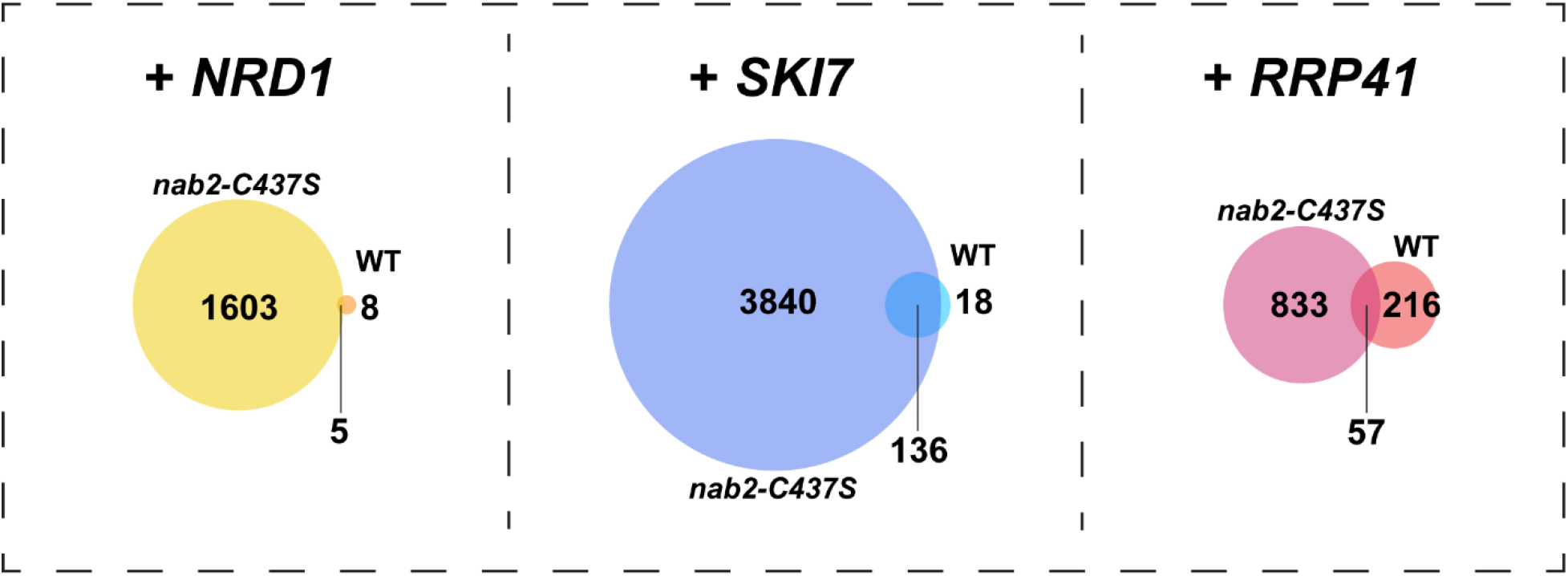
Overexpression of suppressor genes in control cells affects minimal genes compared to overexpression in *nab2-C437S* cells. RNA-sequencing was performed on control cells overexpressing suppressor genes. Differential expression analysis was performed on samples compared to control cells overexpressed with empty vector. Gene counts include transcripts with both up and down steady-state levels. Absolute log_2_ fold change cut-off set at α>0.5. Statistical significance set at p<0.05.

## Discussion

In a previously described screen (17), we identified specific RNA exosome subunits and cofactors that suppress the cold-sensitive growth phenotype seen in *nab2-C437S* mutant cells. Rrp41 and Rrp42, two structural components of the core RNA exosome, were identified as suppressors. Analysis of the mechanism of suppression suggests that overexpression of these individual subunits of the complex suppress by decreasing RNA exosome activity, potentially due to destabilizing complex formation by disrupting subunit stoichiometry. Further supporting the hypothesis of suppression by RNA exosome impairment is the discovery of suppression of *nab2-C437S* cold-sensitive growth by structural RNA exosome mutants that impede transcript progression through the central channel of the complex.

Impairment of the RNA exosome can interrupt numerous functions of the complex and impede processing of various classes of RNAs. In the nucleus and nucleolus, the RNA exosome largely serves to process non-coding RNAs, including pre-rRNAs, tRNAs, and snoRNAs. It also functions in turnover of RNAs and degradation of misprocessed pre-mRNAs in the nucleus (19,57,58). In the cytoplasm, the RNA exosome functions in mRNA turnover and in the quality control pathway responsible for degrading aberrant mRNAs resulting from non-stop decay, nonsense-mediated decay, and no-go decay (35,59,60).

The diversity of roles performed by the RNA exosome among such a wide array of potential RNA targets requires careful coordination of recruitment and activities. This coordination is facilitated by cofactors. Just as there are diverse roles performed by the RNA exosome, cofactors modulate RNA exosome function through a variety of mechanisms. The specificity of exosome cofactors that also suppress the cold-sensitive growth phenotype of *nab2-C437S* cells upon overexpression may also distinguish particular functions of the exosome that are critical for Nab2 function, as well as Nab2 target RNAs that are crucial for growth.

The isolation of Nrd1, Ski7, Nop8, and Mtr4 as the RNA exosome cofactors able to suppress the *nab2-C437S* growth defect upon overexpression implicates their interactions with the RNA exosome as particularly important for the interplay of the complex with Nab2 (Figure 10). Intriguingly, these cofactors are nuclear, cytoplasmic, nucleolar, and nuclear, respectively, and they function in distinct cellular contexts.

**Figure 10:**
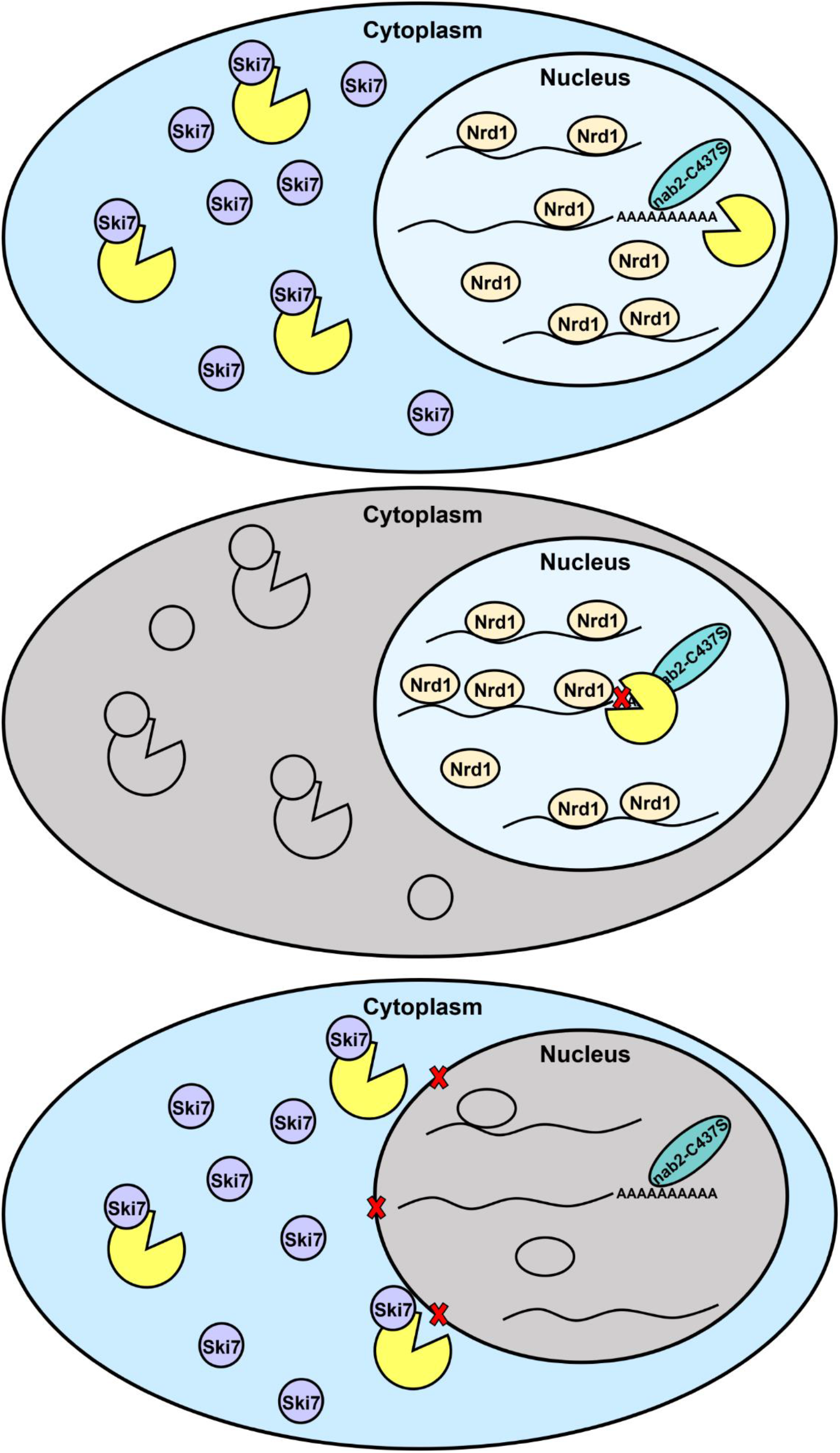
Model: Overexpression of RNA exosome cofactors *NRD1* and *SKI7* impair nuclear RNA exosome function. Weakened binding between Nab2-C437S and target RNAs may result in misprocessed RNAs that are quickly degraded by the RNA exosome. Upon overexpression of *NRD1*, Nab2-target RNAs may be protected from degradation by the RNA exosome due to promiscuous binding of Nrd1. Upon expression of SKI7, the RNA exosome may be sequestered in the cytoplasm, and Nab2-target RNAs may be protected from degradation due to decreased nuclear localization of the RNA exosome.

Although Nop8 was not the focus of this study, its identification as a suppressor does provide supporting evidence for the hypothesis that impairment rather than enhancement of RNA exosome function is responsible for suppression. Nop8 is an essential nucleolar RNA exosome cofactor that negatively regulates activity of the RNA exosome by binding Rrp6 in a concentration-dependent manner. Nop8 functions in opposition to RNA exosome activator Nop53 in the processing of 5.8S pre-rRNA and, ultimately, biogenesis of the 60S ribosome. Nop8 contains an N-terminal domain where it interacts with 5.8S rRNA and a coiled C-terminal domain where it interacts with Nip7, a nucleolar protein which physically interacts with core exosome subunit Rrp43, and Rrp6 (40). As RNA exosome activity is negatively regulated by increasing expression of Nop8, the overexpression of *NOP8* in *nab2-C437S* cells is likely also down-regulating exosome activity.

To further investigate the effect of *nab2* mutation and subsequent overexpression of suppressors, we performed RNA sequencing analysis. We identified fourteen gene ontology categories negatively impacted by the *nab2* mutation. While these categories are largely rescued by *NAB2* overexpression, they are not rescued by overexpression of the suppressors. This suggests the suppressors rescue growth by other mechanisms. Gene sets affected by overexpression of *NRD1*, *SKI7*, and *RRP41* show a large percentage of overlap. This overlap may indicate a common mechanism of suppression, which is likely linked to regulation of the expression of ribosomal subunit genes.

While studying the impact of RNA exosome cofactor and subunit overexpression, a concern arose that our findings could be independent of Nab2. By performing RNA-seq analysis on control cells overexpressing *NRD1*, *SKI7*, or *RRP41* (Figure 9), we found that overexpression of suppressors outside of the impaired Nab2 context had little effect on the transcriptome when compared to control cells without suppressors. Furthermore, we found very little overlap between transcripts affected by suppressors in *nab2-C437S* cells versus control cells. The minimal amount of overlap between the *nab2-C437S* and control cells overexpressing each of these suppressors suggests an important interplay between Nab2-C437S and the RNA exosome. This finding provides evidence that the transcriptomic data acquired is relevant to elucidating Nab2 function and characterizing the functional interactions of Nab2 with Nrd1, Ski7, and Rrp41.

The RNA exosome is localized to both the nucleus and the cytoplasm, but its function is required in the nucleus where it serves as the primary degradation pathway. Conversely, in the cytoplasm, the main degradation pathway utilizes the 5’ to 3’ exonuclease Xrn1 (61). While the RNA exosome functions in both compartments, Nrd1 and Ski7 localize to the nucleus and cytoplasm, respectively. Nuclear-localized Nrd1 assists the RNA exosome in processing nuclear transcripts, and overexpression of *NRD1* likely affects nuclear transcripts. Intriguingly, most transcripts affected by overexpression of *RRP41* are also affected by *NRD1* overexpression, suggesting a primarily nuclear mechanism for suppression by *RRP41*.

Localized to the cytoplasm, Ski7 affects RNAs after they are transported from the nucleus. This may explain the additional transcripts identified as increased or decreased by *SKI7* overexpression. Overexpression of cytoplasmic *SKI7* may also affect nuclear transcripts by impacting the formation or import of the RNA exosome into the nucleus. Through its N-terminal region, Ski7 binds to the same subunits of the RNA exosome as nuclear subunit/cofactor Rrp6 binds through its C-terminal region. These exosome cofactors include Csl4, Mtr3, and Rrp43. Ski7 and Rrp6 bind the exosome in a mutually exclusive manner, and in vitro work has shown that Ski7 can outcompete Rrp6 for exosome binding (21, 62). Overexpression of *SKI7* could impact RNA exosome function by outcompeting Rrp6 and sequestering either binding partner subunits of the exosome or the entire complex in the cytoplasm. This could result in reduced import of the RNA exosome into the nucleus or reduced nuclear localization of binding partner subunits. Either possibility could result in reduced nuclear function of the RNA exosome. Additionally, cytoplasmic sequestration of the complex could lead to enhanced function of the RNA exosome in the cytoplasm by increasing the number of complexes in that compartment. Following up on these hypotheses and elucidating the molecular mechanisms of suppression will be the focus of future studies.

An intriguing possibility is that connections of RNA exosome cofactors Nrd1, Ski7, Nop8, and Mtr4 with exosome cofactor/subunit Rrp6 may provide a common underlying pathway to suppression. Each of these cofactors interacts with Rrp6 either directly or competitively. Nrd1 and Rrp6 work together in the NNS transcription termination pathway (63), as well as a quality control pathway for targeting misprocessed mRNAs for degradation (48). Additionally, Nrd1 may protect transcripts from degradation by Rrp6 by preventing Rrp6 procession along Nrd1-bound RNAs (49). Ski7 and Rrp6 bind to the core RNA exosome in a mutually exclusive manner, potentially affecting localization of the complex (21). Nop8 interacts directly with Rrp6 to down-regulate RNA exosome activity in the nucleolus (40). Mtr4 is recruited to the exosome by Rrp47 and directly interacts with Rrp47 and binding partner Rrp6 at the exosome (64). Mtr4 adenylation activity also competes with Rrp6 deadenylation activity to determine how, depending on transcript stability, RNAs are processed or degraded (43). These connections to Rrp6 could impact Rrp6 function, core exosome function, or both.

Overall, this study strengthens a link between Nab2, the RNA exosome and control of the expression of ribosomal subunit genes. This link could help to explain why the Nab2 orthologue ZC3H14 is so critical in the brain as energy conservation is critical in neurons and ribosome production is one of the costliest cellular processes. Further studies will be required to assess whether the class of RNA targets altered in *nab2* mutant cells in budding yeast is also impacted by loss of ZC3H14.

## Acknowledgements

Financial support as follows - NIH MH107305, AG054206 and GM058728 (AHC). We thank members of the Corbett laboratory for their support.

## Notes

The authors declare no competing financial interests

### Competing Interest Statement

The authors have declared no competing interest.

